# Integrative proteogenomics for differential expression and splicing variation in a DM1 mouse model

**DOI:** 10.1101/2021.05.15.443842

**Authors:** Elizaveta M. Solovyeva, Stephan Utzinger, Alexandra Vissières, Joanna Mitchelmore, Erik Ahrné, Erwin Hermes, Tania Poetsch, Marie Ronco, Michael Bidinosti, Claudia Merkl, Fabrizio C. Serluca, James Fessenden, Ulrike Naumann, Hans Voshol, Angelika S. Meyer, Sebastian Hoersch

## Abstract

Dysregulated mRNA splicing is involved in the pathogenesis of many diseases including cancer, neurodegenerative diseases, and muscular dystrophies such as myotonic dystrophy type 1 (DM1). Comprehensive assessment of dysregulated splicing on the transcriptome and proteome level has been methodologically challenging, and thus investigations have often been targeting only few genes.

Here, we performed a large-scale coordinated transcriptomic and proteomic analysis to characterize a DM1 mouse model (HSA^LR^) in comparison to wild-type. Our integrative proteogenomics approach comprised gene- and splicing-level assessments for mRNAs and proteins. It recapitulated many known instances of aberrant mRNA splicing in DM1 and identified new ones. It enabled the design and targeting of splicing-specific peptides and confirmed the translation of known instances of aberrantly spliced disease-related genes (e.g. *Atp2a1, Bin1, Ryr1*), complemented by novel findings (e.g. *Ywhae, Flnc, Svil*). Comparative analysis of large-scale mRNA and protein expression data showed quantitative agreement of differentially expressed genes and splicing patterns between disease and wild-type.

We hence propose this work as a suitable blueprint for a robust and scalable integrative proteogenomic strategy geared towards advancing our understanding of splicing-based disorders. With such a strategy, splicing-based biomarker candidates emerge as an attractive and accessible option, as they can be efficiently asserted on the mRNA and protein level in coordinated fashion.

**Graphical abstract:** 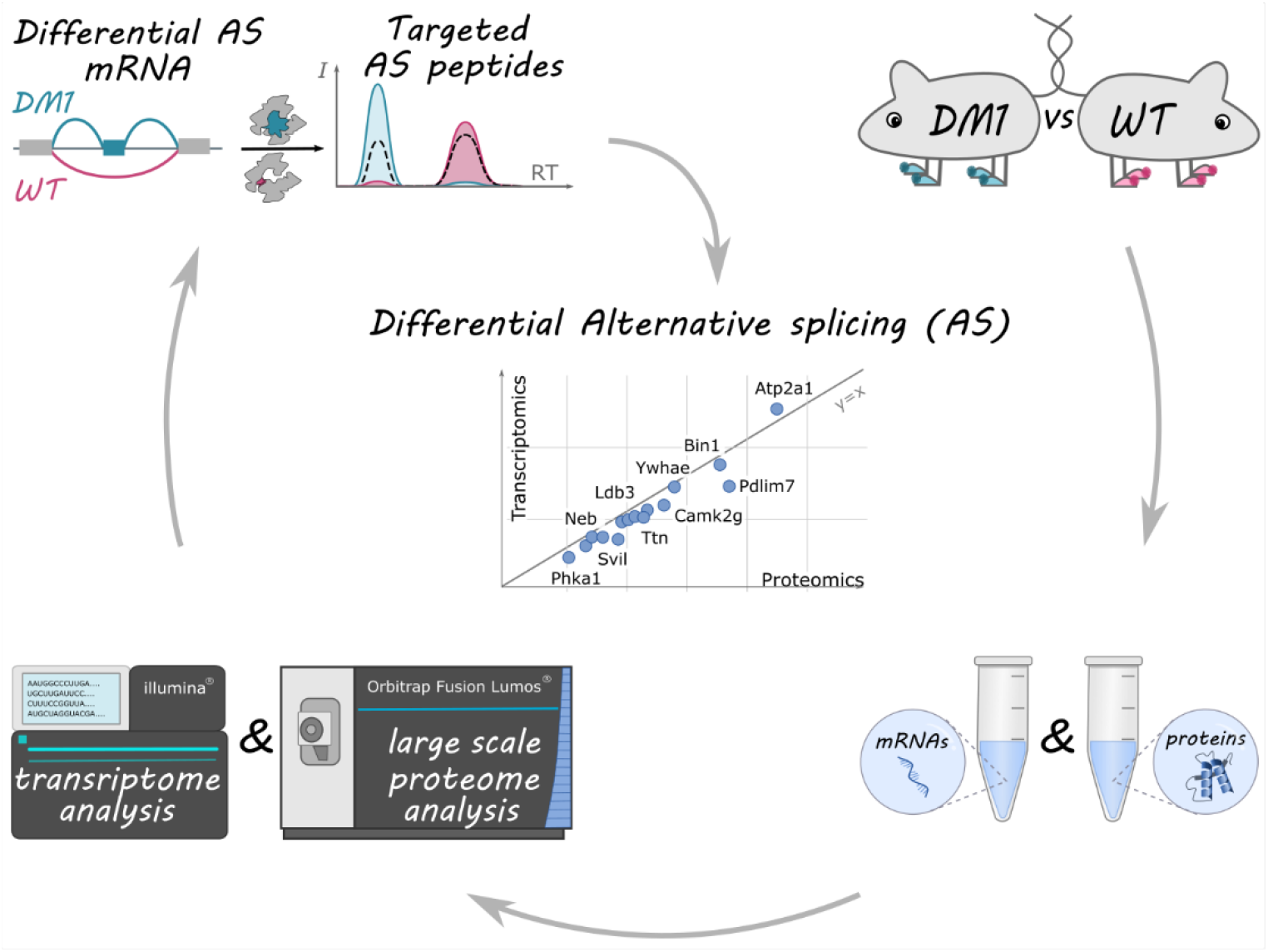

## Introduction

The molecular mechanism of mRNA splicing is essential for creating functional mRNA molecules from disjointly encoded genomic predecessors (exons) in eukaryotes. Alternative splicing (AS) is key for extending, varying, and tuning the arsenal of mRNAs and proteins in the context of developmental or tissue-specific functions. Dysregulation of mRNA splicing, however, is increasingly being recognized as a hallmark of disease, notably cancer (Bonnal et al., 2020), and also degenerative diseases and aging, with interesting implications for shared biological processes (Mateos-Aierdi et al., 2015; Deschênes and Chabot, 2017; Meinke et al., 2018; Raj et al., 2018; Solovyeva et al., 2021). On the transcript and protein level, such dysregulation manifests itself in splicing patterns altered in disease compared to healthy controls, and this differential is measurable with dedicated methods applied to high-throughput transcriptomics (Tx) and proteomics (Px) data. We are hence using the term Differential Alternative Splicing (DAS) to describe instances of altered splicing between two comparison groups.

Modern high-throughput mRNA sequencing (RNA-seq) has become a standard technology for assessing whole-transcriptome gene expression (GE) in biological samples and differential gene expression (DGE). For this work, to extend RNA-seq analyses to instances of altered splicing, or DAS, we specifically employed the LeafCutter software (Li et al., 2018). This tool analyses specifically the exon junction-spanning reads of a genome reference-aligned RNA-seq dataset to identify all sets of splicing alternatives interconnected via one or more shared exons, the so-called “intron clusters”.

Proteome-scale investigations of cellular or tissue-based protein content have become a powerful tool for many biological and medical studies (Aebersold and Mann, 2016). Today, mass spectrometry (MS)-based shotgun Px allows identification of thousands of proteins within several hours (Meier et al., 2018; Demichev et al., 2020). In homogeneous samples from cell lines, the achieved comprehensive coverage of a sample’s protein content is approaching the routine performance of RNA-seq (Bekker-Jensen et al., 2017). However, in complex samples such as tissue, deep coverage is much harder to achieve, as tissue or biofluid proteomes are often dominated by a few highly abundant species. Muscle tissue in particular poses additional challenges due to the complexity and resilience of its tissue architecture (Deshmukh et al., 2015; Drexler et al., 2012).

In MS-based Px, data-dependent acquisition (DDA) is a method where ionized peptides (precursors) are automatically selected and subjected to fragmentation (tandem mass spectrometry). DDA is a powerful discovery modality for profiling a sample’s general protein composition and abundances. However, most of the proteins represented in DDA results are identified by only a few tryptic peptides (Nesvizhskii and Aebersold, 2005). Moreover, the overlap between peptides identified under different instrument settings is often modest due to automatic precursor ion selection. As a consequence, DDA is generally underpowered to systematically investigate a specific set of protein isoforms, although a few methods for alternative splicing analysis at the protein level using DDA were described recently (Komor et al., 2017; Lau et al., 2019; Wu et al., 2019).

As an alternative, targeted approaches (e.g., Parallel Reaction Monitoring, PRM) can reliably quantify peptides with low abundance in a complex mixture. Synthesized analogues of predefined peptides of interest distinguishable via heavy isotope labelling are added to a sample and monitored during its retention time window. These techniques then permit to calculate with high accuracy the ratios of labelled peptides to their unlabelled counterparts naturally present in a sample based on fragment intensities (Picotti and Aebersold, 2012). Targeted approaches have greater sensitivity and reproducibility compared to DDA and allow monitoring of up to a hundred predefined peptides in one run. Moreover, peptide design can be custom-tailored to interrogate specific splice events (Han et al., 2021), informed by sequence database annotation or, as demonstrated here, by adequately processed transcriptomic data.

DM1 (Myotonic Dystrophy, type 1) is a degenerative disease without a cure. It is the most common inherited muscle disorder in adults, with a prevalence commonly reported to vary around 1/8000, (Thornton, 2014), although a recent study ascertained a nearly four-fold higher frequency, pointing to a significant potential for underdiagnosis (Johnson et al., 2021). DM1 is a multisystem disorder that affects skeletal and smooth muscle as well as heart, eye, and the central nervous system. Clinical symptoms involve myotonia (impaired muscle relaxation, a hallmark of DM1), progressive distal muscle weakness and atrophy, cataracts, and cardiac conduction abnormalities. DM1 results from a trinucleotide (CTG) repeat expansion in the *DMPK* gene that is transcribed into RNA, with disease severity positively correlating with the length of the repeats. The excessive non-coding CUG repeats in the *DMPK* RNA bind proteins important for RNA splicing, including alternative splicing. The resulting patterns of splicing abnormalities have been detected in many transcripts coding for proteins important for muscle function (Muge Kuyumcu-Martinez and Cooper, 2006; Nakamori et al., 2013; López-Martínez et al., 2020), among them chloride channels like CLCN1 (Charlet-B. et al., 2002; Mankodi et al., 2002) and calcium channels and pumps like RYR1 (Kimura et al., 2005), ATP2A1 (Hino et al., 2007), and CACNA1S (Tang et al., 2012).

The significant unmet medical need presented by DM1 underscores the importance of a thorough understanding of animal models used to study this disease. The Tg HSA^LR^ mouse model used in our study is characterized by a human *ACTA1* (human skeletal actin) transgene with more than 250 CTG repeats (Mankodi et al., 2000). Post-transcriptionally, the RNA accumulates in the nucleus, where the CUG repeats form extended hairpin structures, which sequester alternative splicing modulators from the ‘muscleblind’ (*Mbnl)* family (Miller et al., 2000), impacting RNA splicing patterns (Ho et al., 2004; Du et al., 2010; Angelbello et al., 2019). Similar to a more recently described inducible DM1 mouse model (Morriss et al., 2018), the CUG repeat expansion is restricted to skeletal muscle. However, unlike human DM1 disease in this model the repeats are not associated with the *DMPK* gene. A better understanding of the splicing alterations in this DM1 mouse model and, ultimately, their similarities and differences to human disease are hence of prime interest.

To date, mostly only isolated individual proteins and protein complexes have been investigated in the context of DM1 disease (Furling et al., 2003; Forner et al., 2010; Hernández-Hernández et al., 2013; Nakamura et al., 2016) with some exceptions specifically focused on the global proteome changes in neurological context (Sicot et al., 2017; González-Barriga et al., 2021). This leaves a notable gap in our understanding of protein and specifically protein isoform changes in this pathological condition. Although the genetic cause of DM1 is known, a better understanding of the molecular disease processes in human and animal model systems is urgently needed to aid progress towards a cure.

To obtain a comprehensive picture of global gene expression in a biological system, the integration of available genomic, transcriptomic, and proteomic data is key (Nesvizhskii, 2014), reflecting an evolved meaning of the term “proteogenomics” since its original introduction (Jaffe et al., 2004). In this work, using an *in vivo* murine DM1 disease model, we describe in detail a highly integrative transcriptomics and proteomics analysis with dedicated focus on differential alternative splicing. Moving beyond simplistic assumptions, we demonstrate that the correspondence between the two technologies is good or even excellent, given the proper experimental constraints and statistical analyses. Our proteogenomic approach not only robustly recapitulates well known splicing changes in DM1, but also identifies and confirms novel ones. It thus provides a viable path to establishing robust splicing-based transcript- and protein-level biomarker and drug target candidates in an integrative setting and is easily generalizable an actionable blueprint for investigations into other splicing-related diseases.

## Materials and Methods

### Animals

Animal experimental procedures in this study conformed to the regulations effective in the Canton of Basel-City, Switzerland, under the license number BS-2885.

Transgenic mice in HSA^LR^ line 20b were maintained as homozygotes on an FVB inbred background (Mankodi et al., 2000), supplied by Novartis Pharma AG, mouse colony MB828. FVB mice were used as wild-type animals and were purchased from Janvier Labs (Le Genest-Saint-Isle, France). The mice were housed at 25 °C with a 12:12 h light-dark cycle in groups of 2 to 4 animals and acclimated to the facility for 7 days. Food and water were provided *ad libitum*.

The right and left hind limb gastrocnemius muscles of 5 HSA^LR^ and 5 WT, 10-12 week-old male mice were dissected, snap-frozen, and stored at −80°C until RNA and protein extraction.

### RNA extraction and short-read RNA-seq

For total RNA, muscles were pulverized, and 15-20 mg were used to extract RNA using a combination of Trizol and RNeasy Micro Kit (Qiagen), followed by a DNase removal step according to the manufacturer. Next generation sequencing libraries were prepared with the TruSeq Stranded mRNA Sample Preparation kit (Illumina) from 350 ng of input RNA using IDT for Illumina TruSeq UD Indexes (IDT). Adapter-ligated fragments were amplified using 12 rounds of polymerase chain reaction (PCR). The resulting libraries were pooled and loaded on two lanes of a HiSeq 4000 (Illumina) for paired-end 76 bp sequencing, generating 71 million reads per sample on average. Reads were aligned for each sample to mouse genomic reference GRCm38/mm10 using STAR v2.5.2a (Dobin et al., 2013) with success rates between 90% and 95%, resulting in 66 million mapped reads per sample on average.

### Bioinformatics: short-read RNA-seq

Analysis of alignment .bam files on the gene level was performed in R/Bioconductor, using the edgeR (Robinson et al., 2010) and limma (Ritchie et al., 2015) packages. The tables ReadsPerGene.out.tab generated by STAR were filtered before statistical analysis, so that only genes with CPM value more than 1 in at least three samples were considered as expressed. Differential expression analysis was performed via linear modelling, accounting for mean-variance relationship. The p-values were adjusted using Benjamini-Hochberg correction for multiple testing.

Analysis of alignment .bam files for DAS was performed using LeafCutter v0.2.7 (Li et al., 2018), requiring at least 20 reads per cluster (-m 20), a maximum intron length of 500 kb (-l 500000), and maximum level of consistency within groups (-i 5 -g 5). Intron clusters were annotated for gene of origin and junction novelty status based on a RefSeq transcriptome (release 90) representing 36 000 annotated genes and more than 100 000 transcripts.

### Proteomics

#### Tissue lysis and digestion, protein extraction

In the light of known challenges to effective protein extraction in skeletal muscle tissue (Gonzalez-Freire et al., 2017), we designed the following extraction protocol. Laboratory chemicals in analytical quality were obtained from Sigma unless stated otherwise. Aliquots of 10 mg frozen muscle tissue powder were resuspended in 250 μL of 50 mM Tris buffer pH 8.5, containing 1% of sodium dodecyl sulfate. Tissue lysis was performed by tip sonication with a Branson Digital Sonifier for 15 sec at 25% amplitude on ice for 3-5 times, with 1 min breaks to allow sample cooling. Proteins were then precipitated using chloroform-methanol extraction (Wessel and Flügge, 1984). After precipitation, protein pellets were dried in a stream of N2 and dissolved in 200 μl 50 mM Tris buffer 8.5 pH with 8M Urea using a sonication bath (Sonorex Digital 10P) (2 x 3 min). Protein extracts were reduced in 5 mM dithiothreitol at 56°C for 40 min and alkylated in 15 mM iodoacetamide (IAA) at room temperature for 30 min in the dark. Finally, to quench IAA excess and prevent overalkylation, DTT was added to the solution to 15mM concentration. For digestion, samples were diluted 3 times with 50mM Tris buffer (down to 2M final urea concentration) and digested overnight at 37°C using a Trypsin/Lys-C mixture (Promega) at a ratio 1:100 w/w. To reduce the number of missed cleavages, an additional digestion step was performed for 3 hours with the protease mixture at the ratio 1:200 w/w. Enzymatic digestion was terminated by adding trifluoroacetic acid to 1%. After the reaction was stopped, the sample was centrifuged at 10 000×g for 5 min, followed by supernatant cleanup on a C18 Sep-Pak cartridge (Waters) according to manufacturer protocol. Finally, the samples were dried in a SpeedVac (Thermo Scientific). The total peptide concentration was measured by UV absorbance at 210 nm using an LC system with UV detector (Agilent), with roughly 10 mg tissue yielding 500-750 μg peptide mixture.

#### TMT-based proteomics

For TMT-labeling, 100 μg of peptides from each sample were processed according to manufacturer instructions (TMT 10plex, Thermo Scientific). After labelling, all samples were mixed and desalted on C18 Sep-Pak cartridges (Waters) according to manufacturer protocol, followed by dissolving samples in 10 mM ammonium formate buffer for high-pH reverse phase fractionation. An Agilent1200 Series HPLC system with a Waters XBridge column (C18 3.5 μm, 150 x 2.1 mm) was used to fractionate samples into 72 fractions during a one-hour linear gradient from 0% to 50% solvent B (ACN with 20mM ammonium formate pH 10). The 72 fractions were concatenated into 24 fractions, dried, and dissolved to 0.3 μg/μL concentration (2% ACN with 0.1% FA) for further analysis.

LC-MS/MS/MS analysis was performed using an Orbitrap Fusion Lumos mass spectrometer (Thermo Scientific) coupled with a Thermo Easy-nLC system. From each high pH fraction, 5 μL of peptides were injected and separated on a heated (55 °C) Aurora nanoZero C18 column (1.6 μm,75 μm i.d. x 250 mm, 120 Å) (Ion Opticks). Mobile phases were as follows: (A) 0.1% FA in water; (B) 90% ACN, 0.1% FA in water. Peptides were eluted using a linear gradient from 2% B to 6% B for 50 min, followed by a linear gradient to 15% B for 50 min and to 30% B for 68 min at a flow rate of 400 nL/min. The column was washed at 95% B for 20 minutes and equilibrated to the start concentration of mobile phase B. Mass spectrometry measurements were performed using data-dependent acquisition (DDA) mode (Top Speed, 3s/cycle), using an MS3 method for accurate TMT quantification (Ting et al., 2011).

MS1 scans were measured in the Orbitrap mass-analyzer with following settings: mass range from 350 m/z to 1,500 m/z, resolving power of 120K, maximum injection time (IT) set to 50 ms, automatic gain control (AGC) for MS1 was 2.0e5, dynamic exclusion set to 40 s. Precursor ions were isolated using a 0.7 Th window, followed by fragmentation using collision-induced dissociation (CID) at normalized collision energy (NCE) of 35%. Fragment ions were measured in the ion trap with maximum IT of 50 ms and AGC value of 1.0e4. After MS/MS, 5 notches were isolated with 2 Th window for high-collision dissociation (HCD) at NCE 55%. The MS3 scans were measured in the Orbitrap mass-analyzer in the mass range from 100 m/z to 500 m/z with resolving power of 50K, maximum IT 50 ms and AGC value 1.0e5.

#### Peptide design for targeted proteomics

For the target tryptic peptides, we adhered to the following design principles: each intron cluster was targeted by ideally at least one peptide specific to each DAS event (e.g., an exon inclusion and an exon exclusion-specific peptide in case of an alternatively spliced cassette exon). Where impossible, interrogating only one alternative was acceptable. In addition to splice event-specific peptides, one “normalizing peptide” per gene was chosen, which – based on available transcript annotations – is common to all known isoforms and unique to this gene.

For each chosen DAS event, amino acid sequences were translated from RefSeq transcripts representing the event or, for exonic sequences without matching RefSeq transcript, from the genomic sequence in the inferred reading frame. For each gene, a normalizing peptide was chosen according to three rules: – (i) it corresponds to a region common for all known transcripts, (ii) it has a high potential detectability according to Peptide Atlas (Desiere, 2006; Deutsch et al., 2008), and (iii) it has a minimal number of amino acids with possible modifications, such as asparagine and glutamine (deamidation), methionine (oxidation), or known phosphorylation sites. In case of highly probable missed cleavages (close arginine and/or lysine), both peptides were considered. Only peptides with length 6 – 21 amino acids were eligible. Based on LeafCutter DAS results, 30 genes were chosen for parallel reaction monitoring (PRM) analysis. In total 98 SpikeTides L peptides with heavy labelled arginine or lysine were synthesized (JPT, Berlin, Germany) with an approximate amount of 10 nmol per peptide.

#### Targeted proteomics

For PRM analysis, samples were dissolved in 2% acetonitrile (ACN) with 0.1% formic acid (FA), spiked with heavy labelled peptides. 1.5 μg of peptide mixture was injected for analysis. The amount of spiked heavy labelled peptides was adjusted to obtain similar ion current (the final amount varied in the range of 10 – 100 fmol per sample). The analysis was performed using an Orbitrap QExactive HF-X mass spectrometer (Thermo Scientific) coupled with a Thermo Easy-nLC system. For each sample, the analysis was conducted twice using two different chromatographic conditions. Under the first condition, a 90 minute gradient (from 3 to 40 % solvent B) and a two-column set up with a C18 PepMap100 trap-column (5μm, 300μm i.d.x 5 mm, 100Å, Thermo Scientific) and a 25 cm analytical column (EASY-Spray PepMap C18, 2 μm, 75μm i.d, 100Å, Thermo Scientific) were used. This condition is more stable and appropriate for a scheduled PRM analysis with a large number of monitored peptides, but the depth of separation is not sufficient to analyze the low-abundance protein isoforms, especially in the case of muscle tissues. To overcome this obstacle, a 150 min gradient with a longer 50 cm analytical column (EASY-Spray PepMap C18, 2 μm, 75μm i.d, 100Å, Thermo Scientific) was employed as a second condition. Mass spectrometry measurements were performed using targeted PRM mode with the following settings: mass range from 340 m/z to 1,200 m/z, resolving power of 60K, maximum injection time (IT) set to 50 ms, automatic gain control (AGC) for MS1 was 1.0e6, dynamic exclusion set to 50 s. Precursor ions were isolated using a 0.7 Th window followed by their fragmentation using high collision dissociation (HCD) at normalized collision energy (NCE) of 27%. Fragment ions were measured with maximum IT of 100 ms, AGC value of 2.0e5, and resolution 60K.

#### Proteomics data analysis

For TMT-based experiments, database searching was performed using Thermo Proteome Discoverer (version 2.1.0.81) with Sequest HT search engine and Percolator post-processing against a RefSeq mouse database (RefSeq release 91, 77337 entries, 58517 non-redundant protein sequences). Search parameters were as follows: 10 ppm for precursor mass tolerance; 0.6 Da for fragment mass tolerance; maximum 2 missed cleavage sites; fixed carbamidomethylated cysteine, TMT labelled N-terminus and lysine; potential methionine oxidation. The identified peptides were filtered to 1.0% false discovery rate. The threshold for precursor contamination was set to 50%, and the minimal average reporter S/N to 10; all missing values and shared peptides were excluded from quantitation analysis. The TMT-intensities were normalized inside each sample to the total intensity as well as to the average value across the wild type group. The differential abundance analysis was done using moderated t-statistics as implemented in the R package applying limma test with Benjamini-Hochberg correction for multiple testing. The search against Canonical Uniprot database (25021 entries, 23445 non-redundant protein sequence) was done with the same parameters.

The Skyline software (4.2.0.19009) (MacLean et al., 2010; Pino et al., 2020) was used for PRM analyses. The library was built based on Mascot 2.4 (Matrix Science, London UK) (Perkins et al., 1999) identifications of LC-MS/MS runs with the excess heavy labelled peptides spiked into a mixture of samples. The fragments from the second to the penultimate ion were used for quantification to increase specificity. The identifications with “rdotp” and “dotp” indexes lower than 0.7 and 0.9, respectively, were excluded. All matches with an “rdotp” index lower than 0.9 were manually checked for fragment co-elution and chromatographic peak shape. Only fragments represented in all samples were considered, others were excluded manually. For all further analyses, custom Python scripts were used.

## Results

### Differential gene expression and differential alternative splicing analysis of RNA-seq data

To investigate transcriptomic changes associated with DM1 disease, short-read RNA-seq analysis was performed on two groups: gastrocnemius muscle tissue obtained from (1) five HSA^LR^ mice (DM1 disease model), and (2) five healthy (wild-type) mice. After data processing (see Materials and Methods), 12861 genes were quantified and compared between the two groups using moderated t-statistics as implemented in the R package *limma* (Ritchie et al., 2015). 374 genes showed significant differential expression (DGE) of at least 2-fold and p-values lower than 0.01 after multiple-testing correction (Figures 1B and 2A, Table S1).

**Figure 1.**
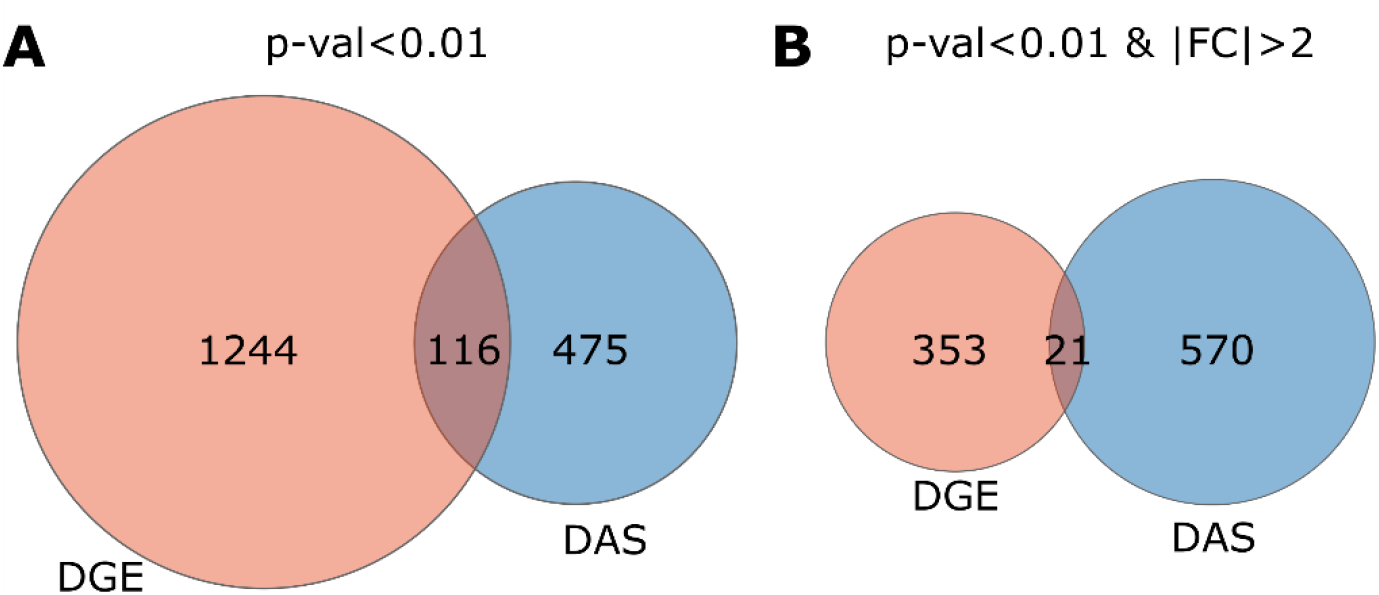
Gene sets showing differential gene expression (DGE) and differential alternative splicing (DAS) and overlaps. (A) Adjusted p-value cutoff p<0.01 for both DGE and DAS. (B) Adjusted p-value cutoff p<0.01 for both DGE and DAS and an additionally for DGE, an absolute abundance fold-change of >2.

**Figure 2.**
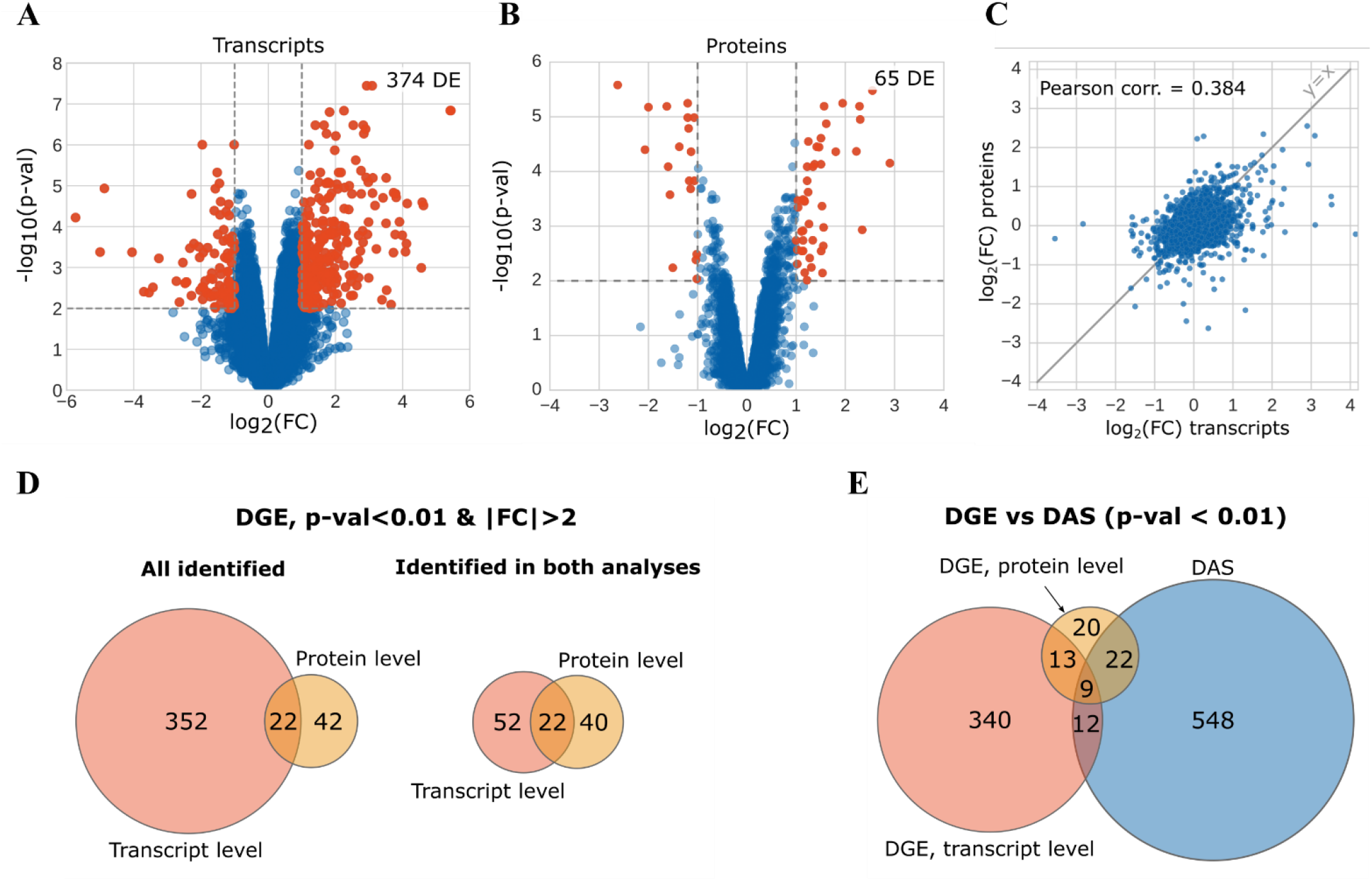
Differential analysis of (A) transcripts and (B) proteins identified in wild-type compared to DM1 mice (volcano plots). The coordinates are log-transformed and correspond to adjusted p-values and abundance fold change calculated by *limma* test with Benjamini-Hochberg correction. The red dots represent differentially expressed transcripts and proteins (p-value < 0.01 and absolute abundance fold change > 2). (C) The correlation between observed transcript and protein abundance fold changes in log-transformed coordinates. (D) The intersection between gene sets with significant differential expression (DGE) on transcript and protein levels (note that 65 DE proteins correspond to 64 genes) and (E) their intersection with genes showing differential alternative splicing (DAS).

We used the LeafCutter software (Li et al., 2018) to identify intron clusters (i.e. sets of splicing alternatives sharing at least one exon). In a second step, LeafCutter statistically evaluates the relative differential usage of the exon junctions (arcs) in each intron cluster between any two comparison groups of interest within the dataset. We equate statistically significant intron clusters with instances of DAS, and we use the term “splice event” to denote a specific splicing variant, or non-circular path of arcs, traversing an intron cluster (see Box 1).

##### Box 1

Three examples of DAS instances shown as LeafCutter-identified intron clusters with different complexity and graphical explanation of terminology in the first example.

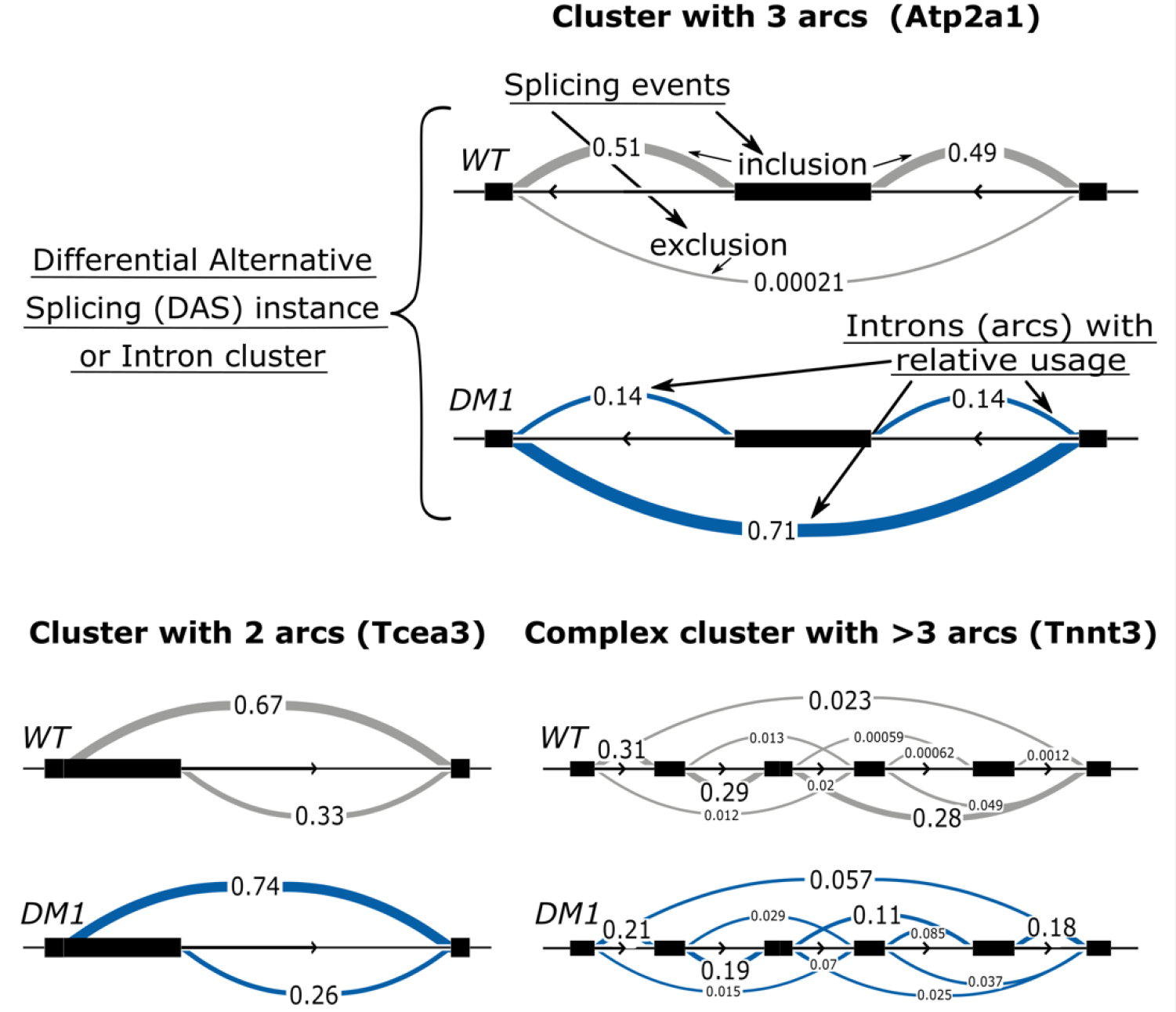

We identified 694 instances of DAS in 591 genes based on a multiple-testing-adjusted p-value cut-off of p < (Table S2). Of these, 268 (39%) intron clusters contain exactly 3 arcs (exon-exon junctions), often corresponding to an alternative in- or exclusion of a single cassette exon flanked by two constitutive exons.

The remaining intron clusters have either only two arcs (typically due to alternative starts or exons with a long and a short alternative, 13%) or a more complex structure (multiple cassette exons and/or variable length exons, 49%). Interestingly, more than half (392) of all identified intron clusters have at least one exon-exon junction not present in the RefSeq transcriptome annotation used. These junctions include unannotated exon pairings or 5’ or 3’ exon boundaries, or combinations. Out of a total of 3161 arcs, 926 (29%) were novel representing splice junctions absent from the RefSeq annotation. Some of these unannotated events reflect simple omissions in the transcriptome reference, or are likely due to errors in the genome reference or sequencing data, but far from all instances can be explained in this fashion. For example, we identified a novel exon in an intron cluster in the *Flnc* gene and its corresponding protein product (see below), with no obvious prior evidence in any of the common annotation sources. We confirmed existence of a novel exon on the protein level for two more genes, *Svil* and *Ywhae,* and a novel junction (exon exclusion) for one more gene, *Ttn* (Table S3). This demonstrates that at least some splicing novelty we observe is real and can be robustly discovered with our approach.

For this work, we focused on intron clusters with 3 arcs, of which 97 (36%) contained at least one unannotated splicing arc. We made this choice because the 2 arc events are enriched for changes that do not alter the coding sequence (alternative transcription starts) and the more complex events increase the challenges for reagent design and data interpretation. Genes with known differential splicing patterns in DM1 constitute an intrinsic positive control in this experiment. For example, our data recapitulates the DM1-specifc exclusion of *Atp2a1* exon 22 (p.adj = 6.48*10^−23^) described for human muscle biopsies from DM1 patients (Hino et al., 2007), for more examples see Figure 4, Discussion and Table 2.

**Table 1.** The number of successful peptides and corresponding genes at different stages of PRM analysis. Number of peptides having “normalizing” peptides represents the total number of peptides used to calculate isoform ratios.

| PRM analysis stage | # peptides | # Corresponding genes / splice events |
| --- | --- | --- |
| Synthesized peptides | 97 | 30 / 45 |
| Heavy peptides detected | 87 | 25 / 40 |
| Splice event-specific peptides detected in samples | 66 | 20 / 27 |
| Normalizing peptides detected in samples | 57 | 16 / 21 |
| Splice event-specific and normalizing peptides, same gene | 47 | 16 / 21 |
| Splice event-specific and normalizing peptides, same gene, in WT and DM1 condition | 37 | 14 / 16 |

**Table 2:**
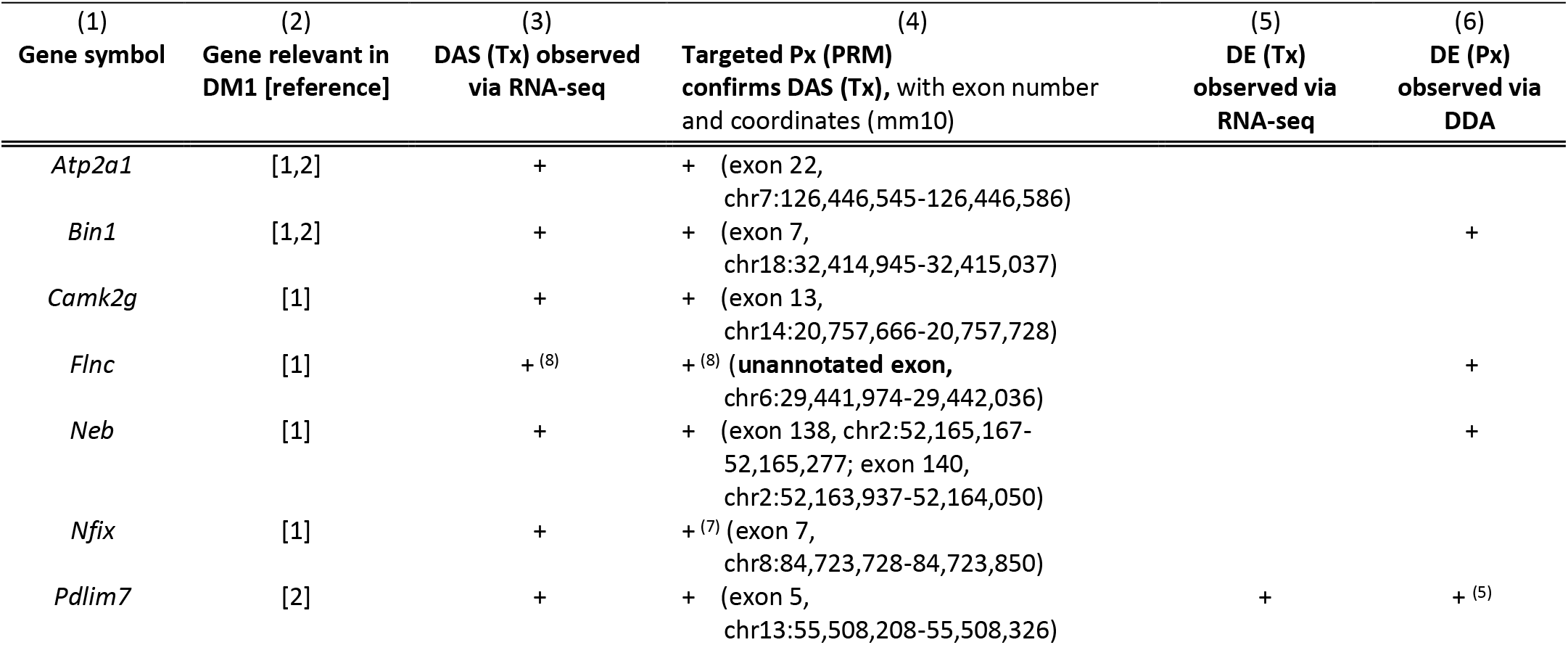
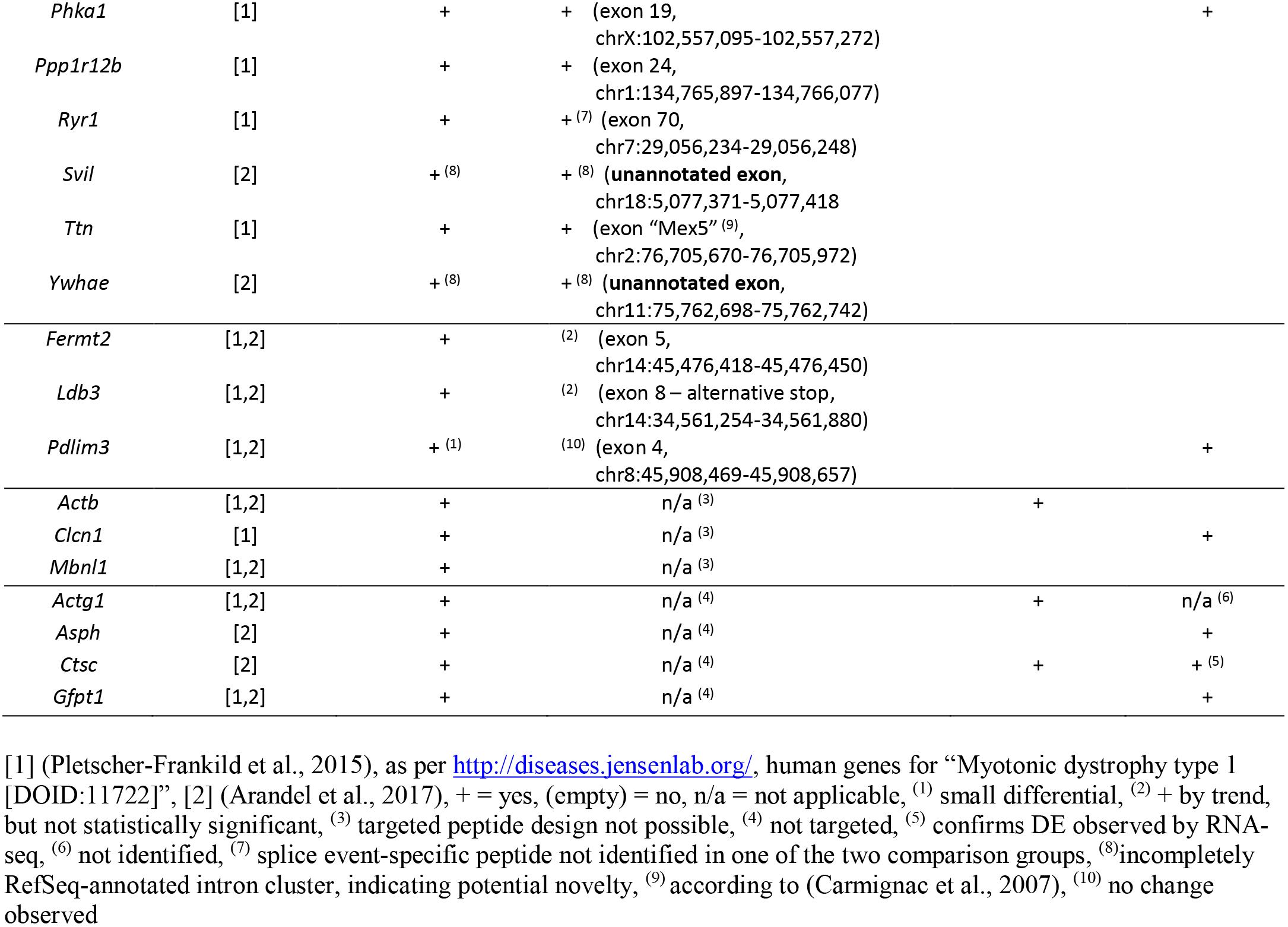
Overview of select genes of known importance to DM1, and summary of their performance in our work, summarized in columns (3) – (6).

**Figure 3.**
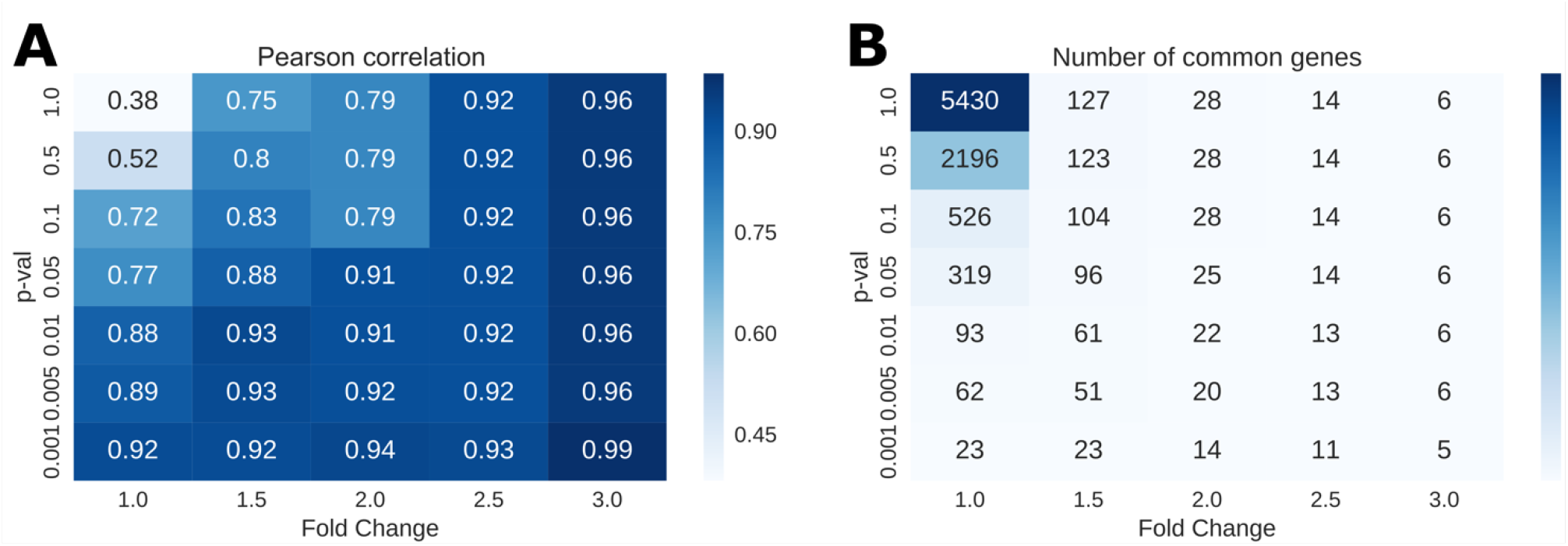
(A) The Pearson correlation of abundance fold changes and (B) the number of common DE genes at transcript and protein levels. Each cell of heat-map corresponds to different significance thresholds, the colour represents (A) the value of Pearson correlation and (B) the number of common genes.

**Figure 4.**
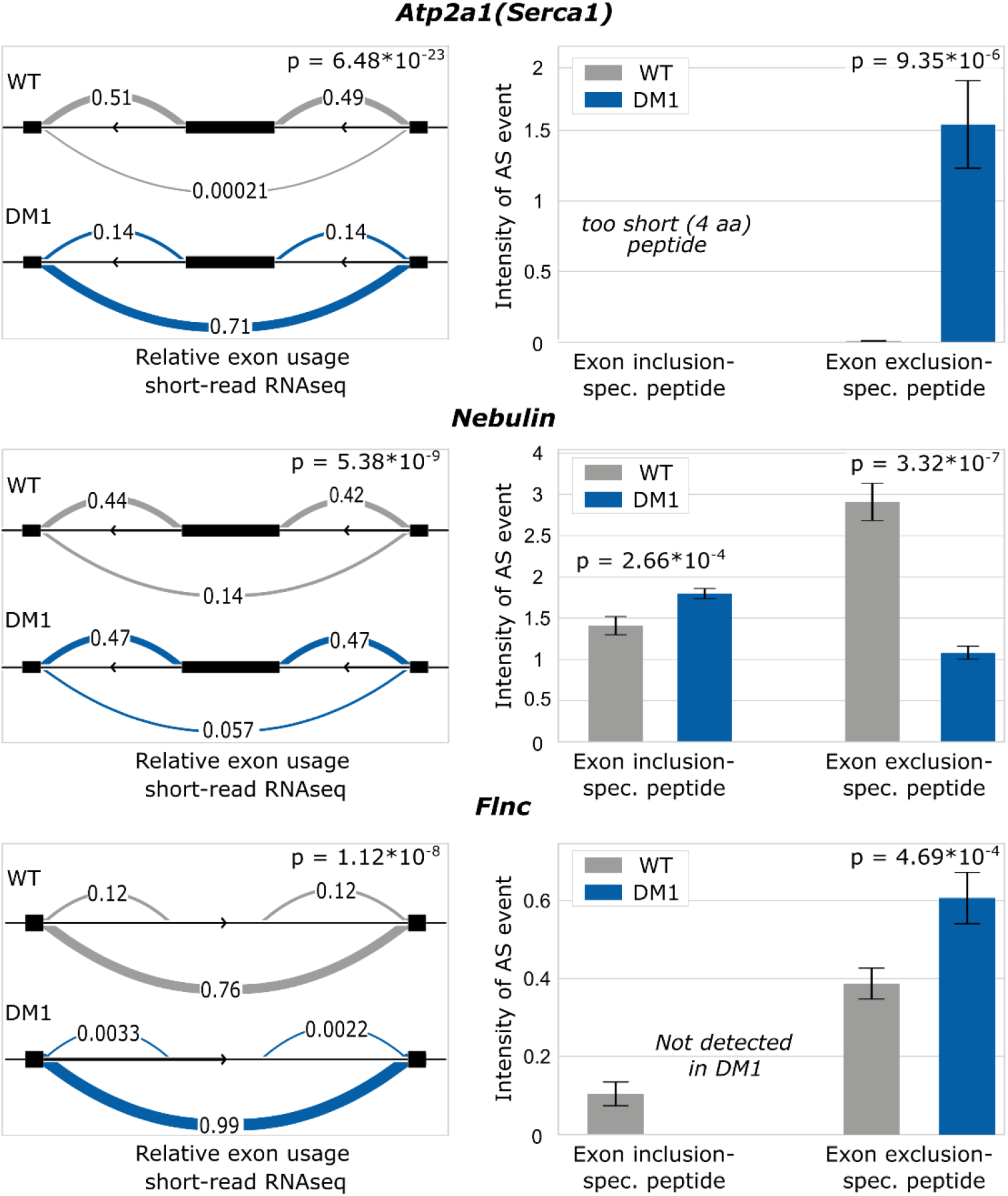
Comparison of instances of differential alternative splicing at protein and transcriptional level for 3 genes (*Atp2a1*, Nebulin and *Flnc*). Panels on the left show LeafCutter AS intron clusters with relative exon usage in the DM1 group (blue) and wild type (WT) control group (grey). Panels on the right show the corresponding relative abundances of peptides specific to exon inclusion and/or exclusion in the same sample groups. See also Figure S5 for a UCSC genome browser-based display of these three LeafCutter AS intron clusters.

We found the overlap between genes exhibiting DGE and DAS to be very modest – fewer than 10% of genes with DGE show DAS, and conversely, fewer than 20% of genes with DAS surface in the DGE analysis (Figure 1A). The dearth of genes in common between DGE and DAS is even more pronounced when imposing a commonly used two fold-change cutoff on DGE (Figure 1B). This reinforces the rationale for separate, dedicated analysis approaches for DGE and DAS, especially in settings like here, where altered splicing is known to be biologically important. We furthermore investigated if these transcriptomic changes actually translate to altered proteins.

### Large-scale DDA proteome analysis; gene expression at protein and transcript levels

To study proteomic changes in DM1 and their correlation with our findings at the transcriptome level, we conducted quantitative MS-based Px analyses on the same ten muscle samples analyzed by RNA-seq. Tryptic peptide mixtures were prepared and labelled using the tandem mass tag (TMT) approach, fractionated to 24 fractions using high-pH reversed phase chromatography and analyzed using high-resolution MS, coupled with high-performance liquid chromatography (HPLC) in a data-dependent manner. To improve the analysis depth, we used extensive prefractionation, addressing the wide dynamic range due to few highly abundant proteins (actin, myosin). Overall, across a total of 24 fractions, more than 53 000 peptides were identified, which resulted in 5832 identified and quantified protein groups.

On the basis of this deep coverage of the muscle tissue proteomes, we identified 65 proteins from 64 genes with significantly different expression between DM1 and control animals (Table S1). Of note, applying the same conventional p-value and abundance fold change thresholds resulted in almost 6 times more differentially expressed genes on the transcriptional level (Figure 2A) than on the protein level (Figure 2B). This considerable difference (summarized in Figure 2D) has likely both technical and biological reasons (see Discussion). Moreover, the overall correlation between DM1-WT abundance fold changes of all genes identified on both the transcriptional and the protein level is low (Pearson correlation = 0.384) (Figure 2C), consistent with general findings from previous proteogenomics studies (Haider and Pal, 2013; Wegler et al., 2020). The intersection between these two sets is modest (Figure 2D): Even if only genes identified in both the Tx and Px experiment are considered (Figure 2D under “Identified in both analyses”), only 30% of genes with DE on the transcriptional level demonstrate DE on the protein level. Moreover, differentially expressed proteins overlap modestly with both DGE and DAS (Figure 2E).

Importantly, the choice of search database (canonical database, Uniprot mouse, 23445 unique sequences vs. full RefSeq mouse, 58517 unique sequences) only minimally impacted the total number of identified proteins. However, 11 of the 31 genes with both DAS and DGE at the protein level were identified only when searching against the full RefSeq database (Figure S1).

The correlation between Tx- and Px-based DGE (0.384 for all genes) markedly increases, when the number of genes is restricted by cutoffs on fold-change and/or p-value, even if these cutoffs are very permissive. For example, applying a p-value threshold of p-val < 0.1 without any fold-change constraints already results in a respectable correlation of 0.72 (Figure 3A) for a gene set 10 times smaller (526) than the total number of genes in common between Tx and Px (5430) (Figure 3B). Imposing commonly used cutoff values (|FC|>2 and p-val < 0.01) yields 22 genes with a correlation of 0.91, and for the five most significantly changing genes, the correlation is almost perfect. This illustrates that the low correlation observed for the complete data set is predominantly determined by noise, while there is substantial agreement between the two technologies for differentially expressed genes beyond standard significance and fold-change cutoffs.

### Design of splice event-specific targeted peptides

For further focused investigation of particular DAS instances and their effect on proteins, we used a targeted approach. We applied parallel reaction monitoring (PRM), allowing precise detection of peptides using heavily labeled synthetic analogues. This method was used to compare the abundance of specific peptides corresponding to parts of proteins predicted to be differentially alternatively spliced between DM1 and WT samples by RNA-seq data analyzed for DAS by LeafCutter. In addition to splice event-specific peptides, one “normalizing peptide” per gene, common to all known isoforms, was chosen. For detailed design principles, see Materials and Methods.

We evaluated 50 simple intron clusters from 40 genes ranked near the top by their LeafCutter-generated DAS p-values. In about 20% of splice events, lysine or arginine residues were present at either the exon start or end, abrogating the possibility of designing a tryptic junction-spanning peptide. Furthermore, the length constraints (see Materials & Methods) eliminated 30% of all candidate peptides checked.

Ultimately, we designed splice event-specific peptides for intron clusters from 30 genes, with four genes containing two separate clusters each. This amounted to 45 distinct splice events, for which 97 heavy peptides were synthesized and used for PRM analysis of the 10 samples (same as used for Tx and Px analyses described above). PRM analysis was conducted twice per sample with different instrument settings (see Materials & Methods). Stages of the analysis were marked by distinct success rates (Table 1), the most prominent obstacle being the detection of the splice event-specific peptides in the biological samples (66 out of 87 peptides). As expected, we observed a correspondence between a gene’s abundance (FPKM values from the RNA-seq analysis) and the recoverability of its "normalizing" and splice event-specific peptides in targeted Px (Figure S2). Altogether, we reliably quantified 21 events in 16 genes. Among these, for 8 events with splice event-specific peptides identified in only one group (WT or DM1), results qualitatively correspond to the Tx results (Table S3 and Figure S3). For 3 of them, there were other peptides covering the same event, which all together sum up to 16 splicing events in 14 genes quantified in both conditions.

### Quantitative analysis of differential alternative splicing in proteins and transcripts

##### Box 2

To quantitatively describe splice events and DAS instances observed between DM1 and WT in our PRM analyses, and to furthermore compare them with their counterparts originating from LeafCutter-processed RNA-seq data, we used the following definitions.

The relative **peptide intensity** in a PRM analysis is calculated as the ratio of the abundances of the light peptide and the heavy (labelled) peptide: *I*^*light*^/*I*^*heavy*^

We then quantified the **relative intensity of a local splice event** (variant) as the ratio between the abundances of the splice event-specific peptide and the normalizing peptide (equation 1),

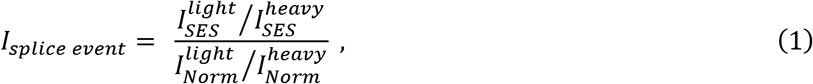

where *I*_*SES*_ is the intensity of the splice event-specific peptide, *I*_*Norm*_ is the intensity of the normalizing peptide, and the indexes light and heavy indicate light and labelled peptides, respectively.

**Protein-based splice event ratios** were calculated as follows for each splice event-specific peptide identified in either group (DM1 and WT):

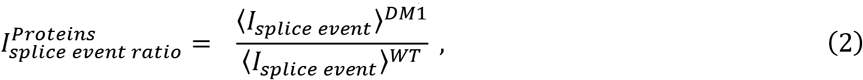

where ⟨⟩ designates the mean value across the group and *I*_*splice event*_ is calculated according to eq.1.

Correspondingly, **mRNA-based splice event ratios** were obtained from our RNA-seq data as follows:

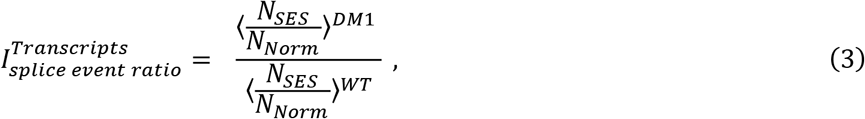

where *N*_*SES*_ and *N*_*Norm*_ stand for the number of reads covering the genome coordinate range corresponding to the equivalent splice event-specific and normalizing peptide, respectively. For exon-exon junction-spanning peptides the number of spliced reads covering this junction was used.

The **relative intensity of a local splicing variant** (eq.1) is similar to the exon usage value calculated by LeafCutter based on short-read RNA-seq data.

A comparison between targeted proteomic results and DAS transcriptomic results shows a good qualitative correspondence. Three examples of targeted splice event intensities calculated according to equation 1 (Box 2) and their corresponding intron clusters from RNA-seq data processed by LeafCutter are shown in Figure 4 and Figure S3, and explained in detail:

1. Skipping of *Atp2a1* exon 22 (based on transcript isoform NM_007504.2), its penultimate exon, is clearly observed near-exclusively in the DM1 condition in short-read RNA-seq data analyzed for DAS and by targeted Px via a peptide-specific to exon 22 exclusion. Unfortunately, it was not possible to design an exon inclusion-specific peptide for this splice event, since it results in an alternative early stop codon and the corresponding peptide contains only 4 amino acids. Importantly, this observation recapitulates findings described in human muscle biopsies from DM1 patients (Zhao et al., 2015).
2. Nebulin (*Neb*) is a gene with more than 150 exons and nearly 30 annotated isoforms in mouse. One of the observed DAS events for *Neb* (intron cluster 5230; see Tables S2 and S3) was targeted by peptides specific for both the inclusion and the exclusion of the alternatively spliced exon. In accordance with LeafCutter results demonstrating the preferred skipping of an exon at genomic position chr2:52,163,937-52,164,050 (mm10) in the WT condition, the peptide specific to this exon’s exclusion has a significantly higher normalized intensity in the WT group. Consistent with this, the exon inclusion-specific peptide has a higher normalized intensity in the DM1 group, and therefore both peptides independently confirm the RNA-seq-based observation.
3. Our data-driven approach enabled us to find novel DAS events, which are not represented in transcript annotations, such as the *Flnc* gene. A cryptic exon (63 nucleotides, “exon 8a” between exons 8 and 9, NM_001081185.2) is included at low levels in WT samples according to transcriptomic data (^~^10% of exon 8 and 9 abundance), whereas its inclusion in the DM1 condition appears negligible (<1% of exon 8 and 9 abundance). Remarkably, we identified the peptide designed to target the 3’ junction of this hypothetical exon in WT samples only. The high correlation between fragmentation spectra of the peptide detected in the sample and the corresponding heavy peptide proves the identification of the targeted peptide, confirming the existence and differential expression of a cryptic exon (Figure S4).

For more examples see Discussion and Table 2.

**Protein- and mRNA-based splice event ratios** (Box 2, equations 2 and 3) resemble the intensity fold changes in a common differential expression analysis, but with a focus on local alternative splicing. Notably, for 8 events, we were not able to calculate this ratio since the corresponding peptides were not detected in one of the groups (e.g. the exon inclusion-specific peptide in the *Flnc* gene was not detected in DM1, Figure 4), but all of them qualitatively agree with the RNA-seq-based results (Figure S3). For 14 genes (16 splice events, with three events targeted by two peptides), splice event ratios were calculated on the protein and transcriptional level (Figure 5).

**Figure 5.**
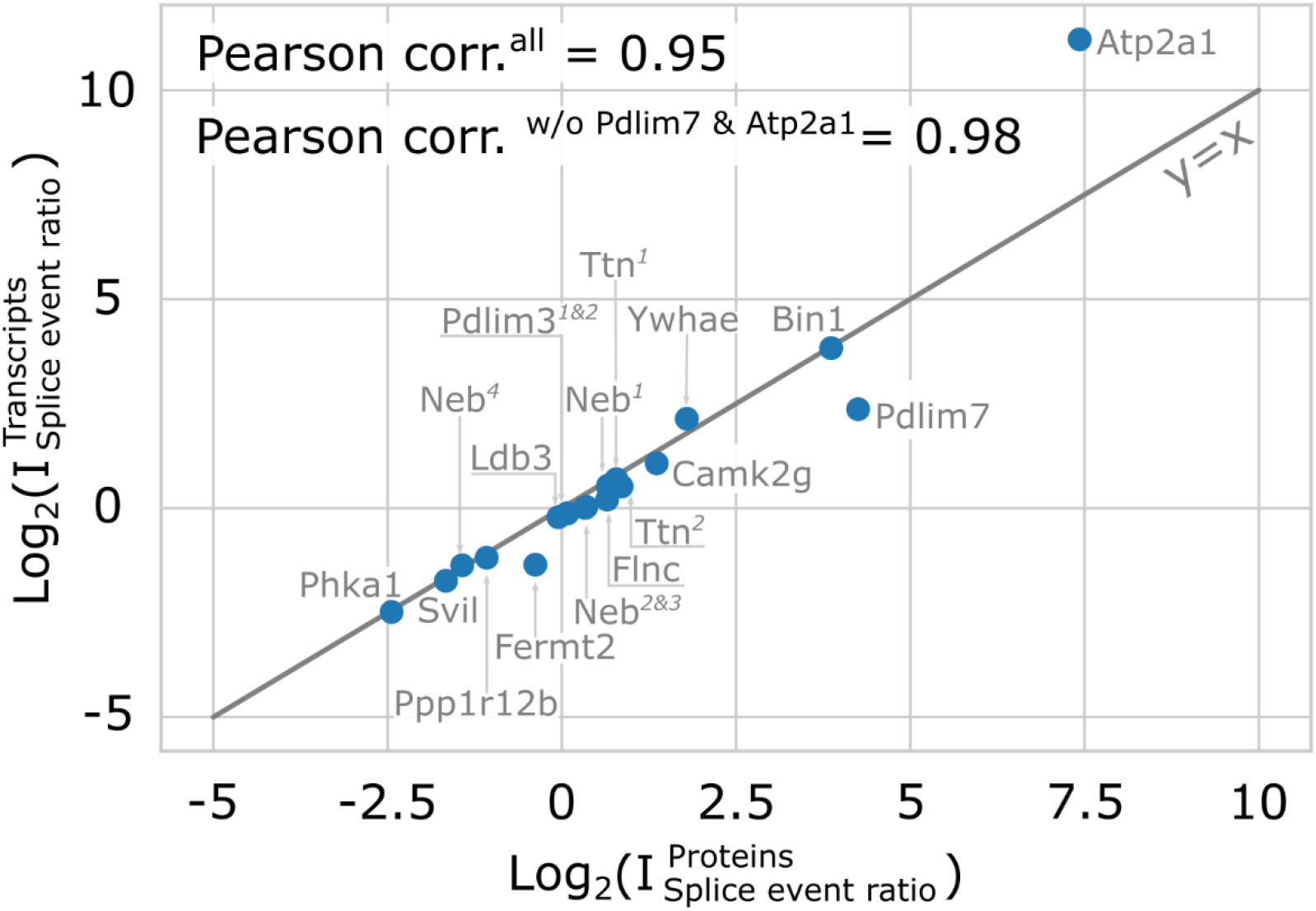
The correlation between log-transformed splice event ratio observed in targeted Px (eq.2, Box 2) and Tx (eq.3, Box 2) for 14 genes, corresponding to 16 splice events and 19 peptide pairs detected in DM1 and WT. The superscript numbers correspond to different clusters/splice events in the same gene (e.g. *Neb*^1^ is an inclusion event in 5321 cluster, *Neb*^2&3^ correspond to different peptides covering the same inclusion event in 5320 cluster, and *Neb*^4^ is an exclusion event in the same 5320 cluster; for all other see Supplemental Table 3 “AS event ratio” list). The Pearson correlation for all examined events is 0.95 (0.98 without two outliers, *Atp2a1* and *Pdlim7*).

The very high correlation (Pearson’s *r* = 0.95) observed between protein and transcriptional splice event ratios demonstrates that DAS events often faithfully translate from mRNA to protein in terms of the relative frequencies of the alternatives in the two sample groups compared (DM1 and WT). *Atp2a1* and *Pdlim7* deviate notably from the diagonal (Figure 5), likely as a consequence of limitations specific to the splice events targeted in these two genes. The transcriptional splice event ratio for *Atp2a1* is the highest (>2000) among all events considered here. But in our targeted Px experiment, the same concentration of heavy peptides was used for all samples across both groups. Therefore, the nonlinear dependency of the peptide ion signal on the concentration may cause inaccurate results for the peptide with the high dynamic concentration range. The *Pdlim7* gene has the most complex of all intron clusters considered in this study: it includes 2 alternatively spliced exons and an alternative length flanking exon, resulting in 7 arcs. This complexity interferes with the ability to accurately calculate the targeted *Pdlim7* splice event intensity. Without these two explicable outliers, the correlation of splice event ratios obtained by Tx and Px is even higher (Pearson’s *r* = 0.98) and demonstrates an outstanding agreement of the results.

We investigated, if the DDA methodology’s generally low peptide identification rate would indeed result in low recovery of the specific peptides targeted in our PRM experiment, as suggested by conventional wisdom. We identified 14 peptides pairs corresponding to 12 targeted DAS events in our DDA data. Moreover, 5 peptide pairs (from 3 genes) were identified exclusively in DDA data. Although our DDA data indicate the known effect of ratio compression in TMT-based experiments (Savitski et al., 2013; Ahrné et al., 2016) compared to PRM data, the correlation between log-transformed splice event ratios for Px DDA and Tx data (Pearson correlation = 0.91, Figure S6B) is close to the one obtained for PRM, and the correlation between the two alternative Px splice event ratios is excellent (Figure S6C). This demonstrates that pursuing even targeted splicing analyses using DDA can be feasible under certain favourable conditions.

## Discussion

In recent years, major advances have been published regarding the transcriptomic landscape of DM1 (Wang et al., 2019) and how alternative mRNA splicing affects the proteome either generally (Liu et al., 2017) or specifically in muscle tissue (Nakka et al., 2018). This work represents, to our knowledge, the first in-depth assessment of a DM1-related *in vivo* proteome based on the HSA^LR^ mouse model, obtained in parallel with transcriptomic data from the same animals. It also provides the first quantitative comparative analysis of splicing changes integrating transcriptional and protein expression data in a DM1-relevant *in vivo* context (Figures 1, 2; Tables S1, S2).

It is important to note that a direct correspondence between transcriptomic and proteomic readouts cannot be universally assumed or expected, for diverse technical and biological reasons. This is true for expression levels and their differentials and can play out on both the gene and the isoform level. An illustrative example presents itself in our data with *Clcn1*, for which we observe significantly lowered expression in DM1 on the protein (Table S1), but not on the transcript level, where however DAS was evident (Figure 2E, ‘DAS’ and ‘DGE, protein level’ intersection, and Figure S7). This observation can be understood based on preferential inclusion of a 79 bp exon in DM1 over WT, leading to a frameshift and premature stop codon. Consequently, these transcripts are subject to elimination via nonsense-mediated decay which abrogates translation, as described before (Charlet-B. et al., 2002).

On this background, it is not surprising that our data indicate an overall limited correspondence between transcriptomic and proteomic gene expression fold change data. However, imposing even modest requirement on the cutoffs for fold change or statistical significance of our two-group comparisons leads to markedly improved correlations for a corresponding subset of genes (Figure 3). This illustrates that one major driver of poor inter-dataset correspondence is noise affecting genes with no actual differential expression. By contrast, differentially expressed genes trend similarly in Tx and Px data, even if much more permissive thresholds for significance or abundance fold changes than commonly used are applied. For the specific instances of DAS identified using both platforms, our data show an excellent correspondence between Tx and Px data (Figures 4, 5, S3; Tables 1, 2; Table S3). This cannot be automatically assumed (see the case of *Clcn1* above), but is generally plausible since the constraints for translating splice events from mRNA to protein are higher than for simple expression level differences, provided the mRNA regions for all splicing events considered are actually protein coding. In this context specifically, our findings also counter the notion that alternative transcripts may generally not result in protein products (Tress et al., 2017) and thus echo the insights from other recent studies designed for isoform detection in proteomic data (Lau et al., 2019).

An important insight emerging from our Px PRM work is that the statistically increased probability of arginine or lysine being encoded at exon boundaries as described in (Wang et al., 2018) is adversely affecting the ability to design tryptic splice event-specific peptides to a noticeable degree in practice. In a setting where choosing a limited number of splicing events from a larger candidate pool is feasible, this limitation may not be overly severe, but it does reduce the generality of such an approach. The problem could be mitigated using enzymes for protein digest other than trypsin, although the available alternatives are less efficient and established (Giansanti et al., 2016).

Following common practice, we used DDA Px initially for establishing the proteomic landscape of differential expression in the DM1-WT context. However, in addition, we also found that analyzing DDA-Px data for splicing events can yield results comparable to targeted approaches. This we ascribe to several key factors. Among these is the choice of a search database that systematically represents splicing variation. Commonly used “canonical Uniprot”-based databases represent annotated splicing variants incompletely or not at all, precluding identification of relevant unique peptides. As we demonstrated using a comprehensive RefSeq-based protein database (Figure S1), systematically including annotated isoforms improves the ability to find splicing-specific peptides without detrimental effects on the total number of identified peptides and protein groups, while keeping the false discovery rate fixed. It also represents a powerful means to improve Tx/Px data integration, as transcripts and their derived proteins are consistently and systematically linked. Other factors contributing to the unexpectedly high success rate recovering targeted DAS-specific peptides in our DDA data are likely the deep sample prefractionation and the long chromatographic gradients we employed, as well as the care we took to optimize sample preparation and mass-spectrometry analyses.

Taken together, these insights suggest an optimized proteogenomics strategy to assess splicing diversity as follows: short-read RNA-seq data are simultaneously analyzed for a gene-level overview and for splicing changes. For these, we show that good tractability and validation rates on the protein level can be expected, when candidate choice is prioritized by high gene expression levels (Figure S2). Px DDA profiling experiments may be conducted and analyzed concurrently with RNA-seq, but ideally before targeted approaches are pursued, so that the former can not only complement, but also inform the latter, in conjunction with the transcriptomic results. Conceivably then, these aggregated and integrated results may guide follow-on long-read RNA-seq of select genes of interest in order to converge on actual full length isoform quantitation. The insights from such long-read data could help constrain possible interpretation of the otherwise local splice event data even on the protein level, where measuring isoforms remains technologically elusive.

It is well known that alternative splicing can be extremely complex, and our transcriptome-wide DAS results reinforce this. Owing to their combinatorial properties, even modestly complex events, involving for example only two alternatively spliced exons, may not yet be conceptually challenging, but are at least burdensome to confirm or track using “local” approaches like qPCR, or, as focused on here, targeted peptides. It is hence important that the complexity of the identified alternative splicing instances can be reliably predicted in the specific context of interest and factored into any prioritization, as exemplified with our RNA-seq DAS analysis approach. In this fashion, the barrier for confirmatory experiments or multi-modal integration (qPCR, Tx / Px) is lowered and chances for success are increased by focusing on simple splicing events, e.g. one alternatively spliced cassette exon flanked by two constitutive exons.

We observed a surprisingly high frequency of intron clusters in our Tx data with one or more unannotated splicing connections, when using a complete RefSeq transcriptome as annotation reference (RefSeq and Ensembl transcriptome annotations are of overall similar size and complexity for mouse) (Table S2). This was unexpected, given that mouse genome assembly and annotations are considered generally mature today, and it illustrates that uncritically relying on existing standard transcriptome annotations is, at this point, still premature. Our cursory investigation showed that many (but not all) novel splicing events have direct or indirect supporting evidence (complementary Ensembl annotations, sequencing data, cross-species sequence conservation) not reflected in RefSeq. Importantly, three of the potentially novel events identified were subjected to targeted Px and confirmed. We are hence illustrating an integrated framework for targeting novel splicing events specifically, starting with their *de novo* discovery using annotation-agnostic RNA-seq data analysis that then constitutes the basis for proteomics-based targeted validation via custom tailored peptide design. It stands to reason that such novel events may be enriched in cases that selectively manifest only in specialized contexts as defined by for example disease, state, stage, or tissue, possibly as lower expression variants. Indeed, the novel *Flnc* variant we found and confirmed was wild-type specific in our comparison and may represent a minor muscle-specific variant lost in DM1 disease.

The high degree of correspondence found underscores the potential of alternative splicing events as robust disease differentiators following the premise that their protein-level manifestation is intrinsically valuable (e.g. for circulating biomarkers) and that establishing such a manifestation is feasible, with a high probability of success, using the *de novo* transcriptomics-guided approach detailed here.

This concept is illustrated in Table 2 below, which shows a selection of DM1-relevant genes with differential alternative splicing, through the lens of the methods and analyses we applied. The first 16 genes in this table are those for which we were able to investigate the transcriptomics-based DAS hypothesis using targeted Px (PRM), fully confirming it in 13 cases and confirming by trend without statistical significance in 2 cases. In one case (*Pdlim3*), the weak but significant DAS event did not correspond to any discernible difference in PRM data.

DM1 muscles show perturbed calcium homeostasis and abnormal excitation-contraction (EC) coupling processes. This involves key regulatory proteins like ATP2A1, CAMK2G, and RYR1, for which we found and confirmed DAS. Several of the differential alternative spliced genes listed in Table 2 code for proteins which play a major role in assembly and function of the sarcomere, the basic contractile unit of the striated muscle fibers (e.g. TTN and NEB), or are involved in sarcomere protein organization (BIN1, LDB3). Yet other proteins (e.g. FLNC, ACTB, SVIL, ACTG1) localize to the Z-disc and are responsible for sarcomere maintenance, integrity and force transduction. FLNC is particularly intriguing due to the novel WT-specific exon we found and confirmed. Given its described essential role for myogenesis, it is tempting to speculate about a narrowing of its functional role under disease conditions. Chloride ion influx stabilizes the electrical charge of the cell, which prevents muscles from contracting abnormally. We recovered described DAS for the *Clcn1* gene (Charlet-B. et al., 2002) encoding the chloride channel 1, which controls the flow of chloride ions into and out of muscle cells and thus plays a key role in repolarization of the muscle after contraction. Our work also confirmed DAS for PPP1R12B, a key regulatory enzyme involved in glycogen metabolism, which supplies the muscle with glucose during contraction. *Nfix* encodes a transcription factor involved in muscle development and regeneration where it controls alternative splicing. MBNL1, another RNA alternative splicing factor known for its prominent role in DM1 pathology and its sequestration in nuclear foci by binding to CUG expansions also shows differential alternative splicing patterns.

Overall, we have demonstrated that differential splicing events between muscles from DM1- and WT mice can be robustly and congruently detected in both transcriptomic and proteomic data in an integrative framework. Furthermore, many of the genes found to be alternatively spliced in this study have been described as playing a role in animal models of DM1 and the human disease itself, or at least have a plausibly relevant function. Generalizing considerations by (Tanner et al., 2021) and extending them from mRNA- to protein-based measurements, we propose DAS events as robust disease biomarker and, possibly, target candidates tractable within the scope of splicing-related disorders such as dystrophies more generally (Pistoni et al., 2010; Scotti and Swanson, 2016), in neurological disease such as Alzheimer’s (Raj et al., 2018), or in the context of aging-related disorders. Indeed, DM1 is considered a disease of premature aging (Mateos-Aierdi et al., 2015; Meinke et al., 2018), and the parallels described between DM1 and aging are strikingly echoed in this work and our related investigations into muscle senescence (Solovyeva et al., 2021). We believe it is remarkable that many of the same genes, with the same splicing events, are affected similarly in an aging model (old rats compared to young) and in the murine disease vs. normal context we describe here. To immediately further these inquiries, validated peptides from this study could be used directly, after species-specific adjustments as needed, for raising splicing event-specific antibodies for diagnostic research.

## Abbreviation list

DM1: Myotonic Dystrophy, type 1
WT: wild-type
Tx: Transcriptomics
Px: Proteomics
GE: Gene Expression
DGE: Differential Gene Expression
DAS: Differential Alternative Splicing
RNA-seq: high-throughput mRNA sequencing
FPKM: Fragments Per Kilobase of transcript per Million mapped reads (RNA-seq analysis)
FC: fold change
MS: Mass Spectrometry
HPLC: High-Performance Liquid Chromatography
DDA: Data-Dependent Acquisition
TMT: Tandem Mass Tag
PRM: Parallel Reaction Monitoring

## Availability

The RNA-seq FASTQ files for this study have been deposited to the European Nucleotide Archive (ENA) via ArrayExpress (https://www.ebi.ac.uk/arrayexpress/) (Athar et al., 2019) under accession number E-MTAB-10842.

The mass spectrometry proteomics data have been deposited to the ProteomeXchange Consortium via the PRIDE (https://www.ebi.ac.uk/pride) (Perez-Riverol et al., 2019) partner repository with the dataset identifier PXD025589.

## Supplementary data

Supplementary Data are available online.

## Acknowledgements

We are grateful to Damien Begue, Peggy Lefeuvre, and Kerstin Oelkers for assisting us with proteomics and transcriptomics data generation, to Yunyu Zhang, Frederique Black, and Gianluca Santarossa for their data analysis help, and to Ulrike Trendelenburg for valuable contributions to the manuscript. We would like to thank Chikwendu Ibebunjo, David Burckhardt, Sophie Dessus-Babus, Robert Bruccholeri, Edward Oakeley, and Oleg Iartchouk for helpful discussions. Finally, we are indebted to Sabine Guth, Estelle Trifilieff, Sophie Lemire, Michaela Kneissel, Karen Wang, Elizabeth Capitelli, Mark Borowsky, and Mikhail Gorshkov for their generous support.

