## Supplemental figures for "Integrative proteogenomics for differential expression and splicing variation in a DM1 mouse model"

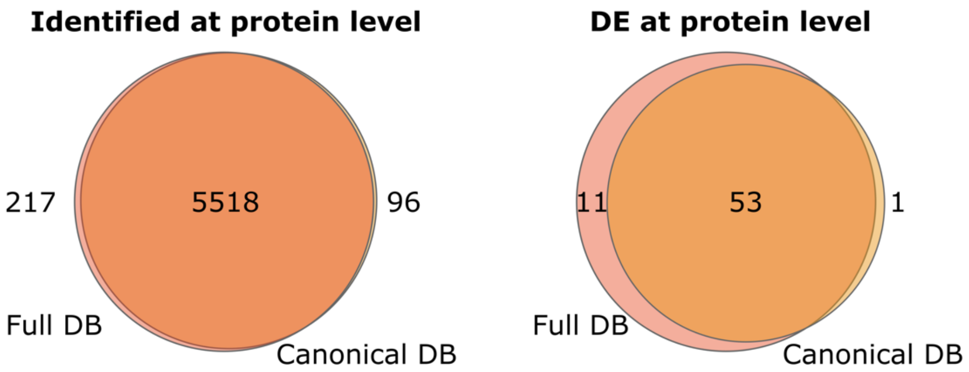


**Figure S1**. The intersection between all **(left panel)** and DE **(right panel)** proteins identified in the search against RefSeq (58517 unique protein sequences) and Canonical Uniprot (23445 unique protein sequences). All DE genes identified only with RefSeq (11 genes) demonstrate DAS as well.

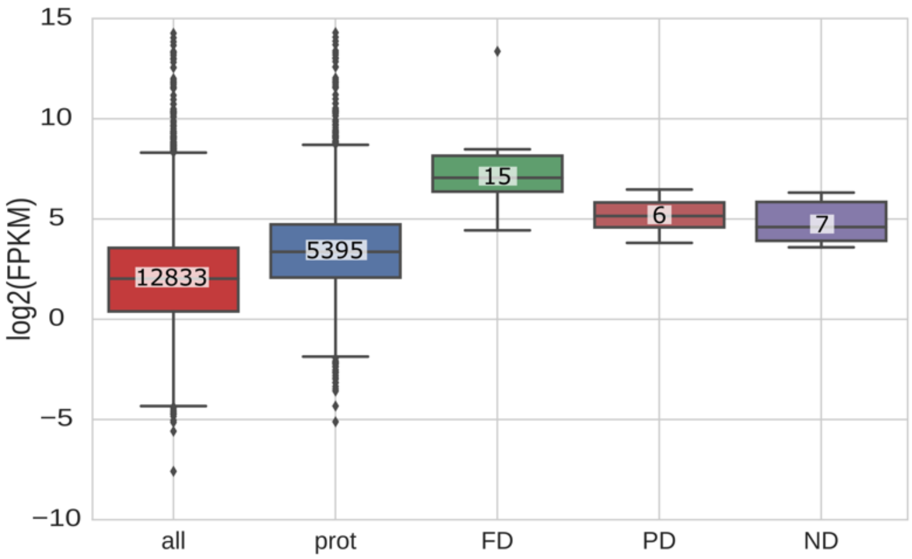


**Figure S2.** The distribution of log-transformed FPKM values of all identified transcripts ("all"), the transcripts with corresponding identified protein in DDA analysis ("prot"), the transcripts for which all peptides were identified in targeted analysis (fully detected- "FD"), the transcripts with only normalizing peptides identified in targeted analysis (partially detected- "PD"), and transcripts with neither normalized nor alternatively spliced peptides identified in targeted analysis (not detected - "ND").

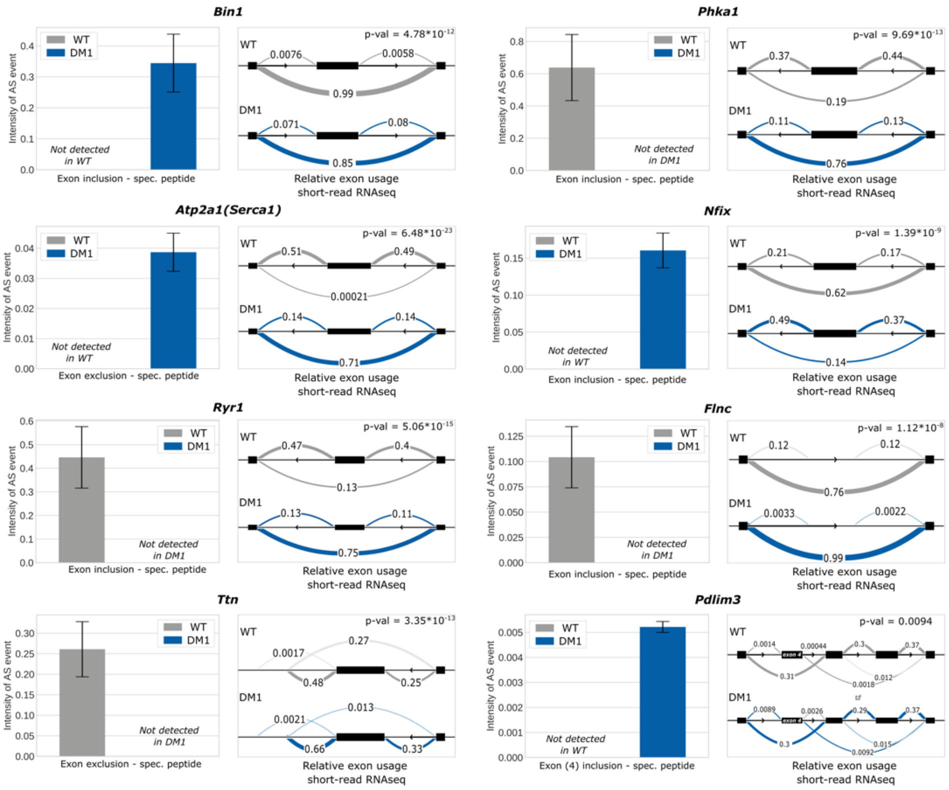


**Figure S3.** Comparison of alternative splicing events at protein and transcriptional level for eight genes with targeted peptides identified in only one group (WT or DM1). The bar plots show relative abundances of peptides specific to exon inclusion or exclusion in the DM1 group (blue) and the wild type (WT) control group (grey). Arc plots show the corresponding LeafCutter intron clusters with relative exon usage in the same sample groups. Here, light grey and blue correspond to 'cryptic arcs'.

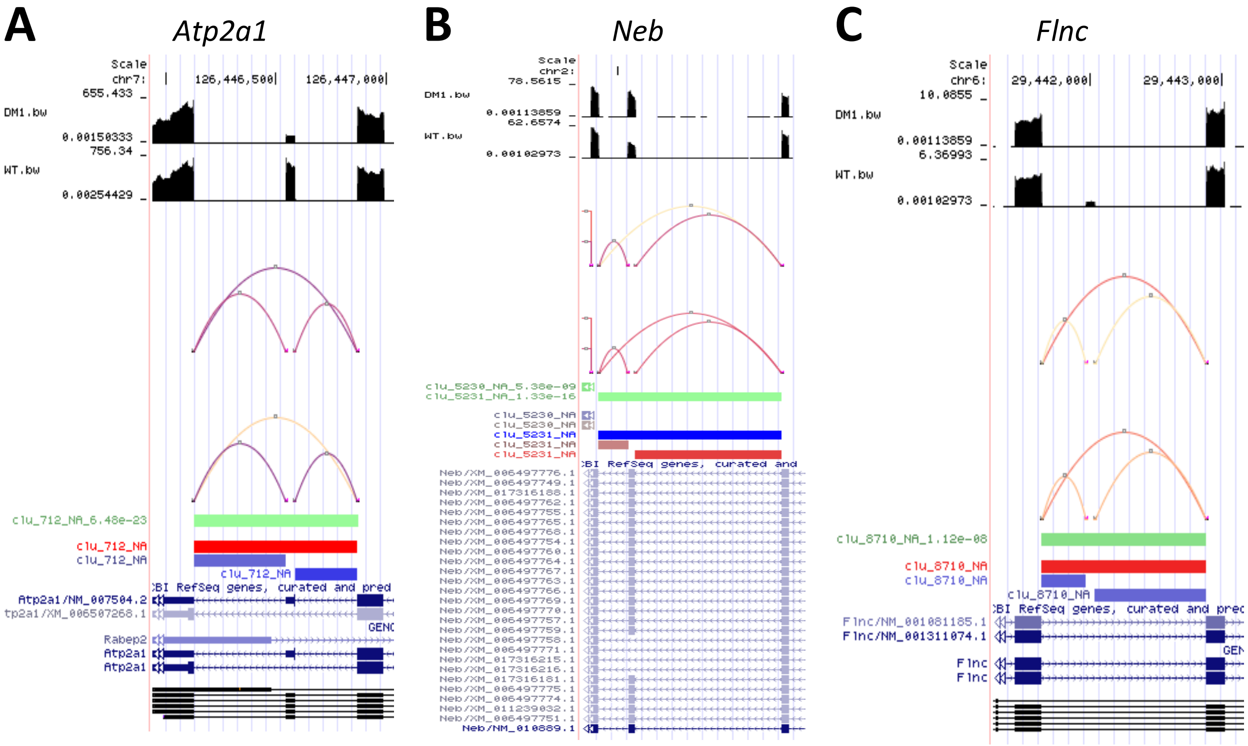


**Figure S4.** Three LeafCutter DAS clusters for **(A)** Atp2a1, **(B)** Neb, and **(C)** Flnc (see also Figure 4). In each panel, data are visualized in three sections, from the top: (i) sample group mRNA expression, (ii) sample group intron junction counts, and (iii) differential splicing between sample groups.

(i) The first pair of bigwig tracks depicts mRNA expression (exon coverage) using group-specific normalized and averaged genome coverage counts. (ii) Intron junction counts for the groups are illustrated with arcs colored according to the "magma" palette**,** with darker hues indicating higher junction read counts, implemented using the UCSC interact data format. Sample-specific raw junction read counts from the LeafCutter clustering algorithm are normalized to mapped read counts and averaged for each sample group. (iii) LeafCutter's differential splicing output is displayed as two bed tracks: a cluster significance track (top) with the genomic footprint of significantly differential intron clusters shaded green and an effect sizes track (bottom), indicating intron start and end positions in shades of red (preferred intron usage in DM1 over WT) and blue (preferred usage in WT over DM1. FDR-adjusted p-values for intron clusters are included in the cluster name.

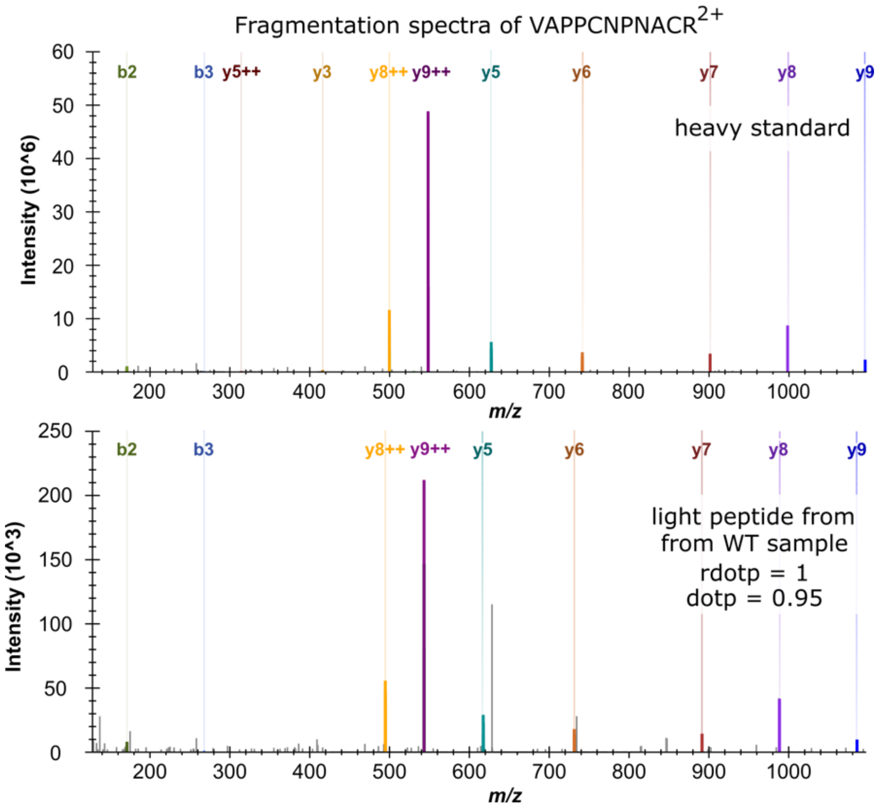


**Figure S5.** The fragmentation spectra of (A) heavy labeled and (B) light peptides observed in one of the wild type mouse samples. The peptide with amino acid sequence VAPPCNPNACR corresponds to the cryptic exon "8a" predicted by LeafCutter in the Flnc gene. The dot product (so-called "rdotp index" in the Skyline software) between the ten most intense fragments of the peptide detected in the sample and the corresponding heavy peptide equals 1 for all WT samples (Skyline "dotp index" varies in the range 0.93 - 0.95)

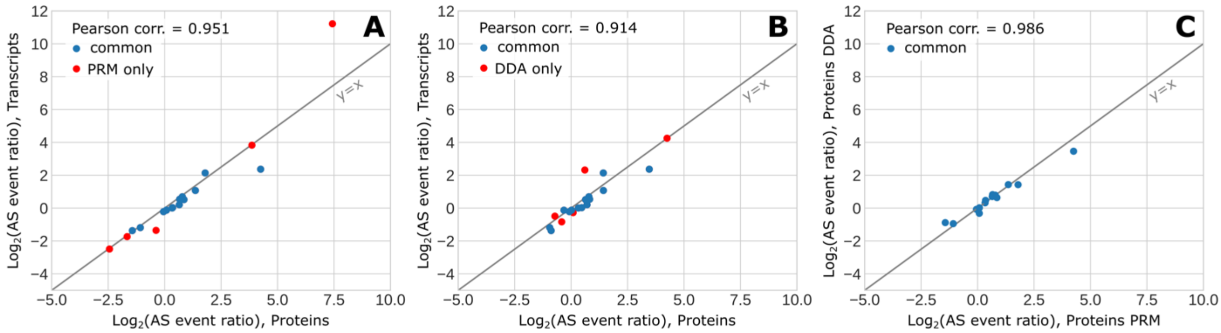


**Figure S6.** The correlation between log-transformed splicing event ratios observed in Tx (eq.3, main text) and Px (eq.2, main text). The intensities of the peptides were obtained from: **(A)** targeted (PRM) and **(B)** DDA analysis. **(C)** The correlation between observed log-transformed splicing event ratios corresponding to peptides identified in both analysis (9 genes, 14 peptide pairs). 5 peptides pairs corresponding to 5 genes were identified only in PRM and 5 peptides pairs corresponding to 3 genes were identified only in DDA analysis.

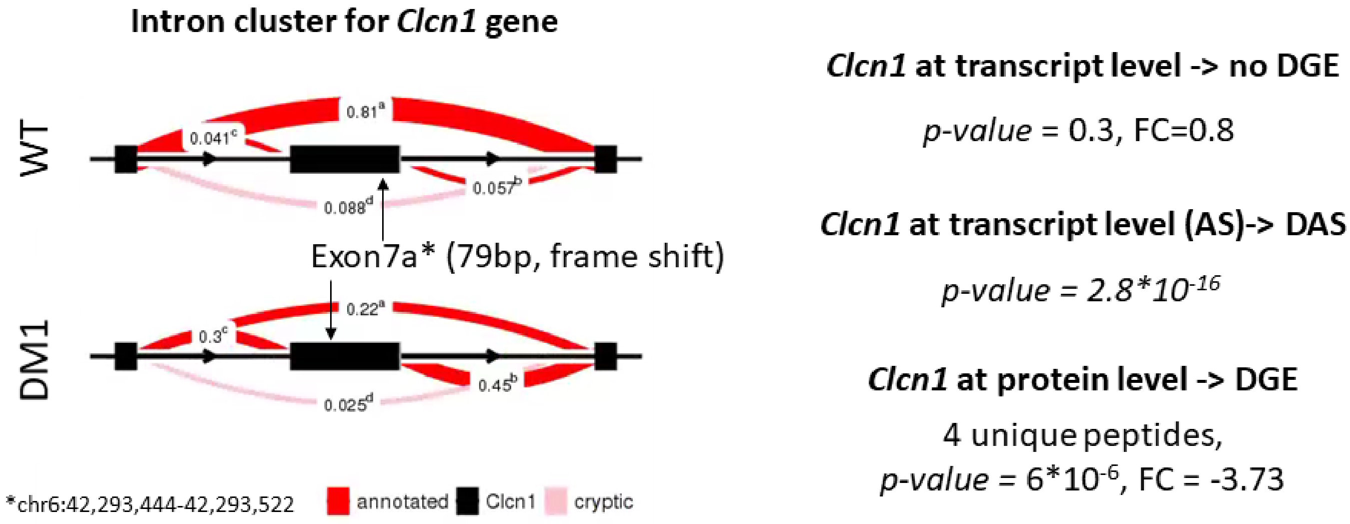


**Figure S7.** The intron cluster identified for *Clcn1* gene (left). The statistics (p-values and fold changes) for the *Clcn1* gene at the transcript and protein level (right).
